# Common variants of *NRXN1, LRP1B* and *RORA* are associated with increased ventricular volumes in psychosis - GWAS findings from the B-SNIP deep phenotyping study

**DOI:** 10.1101/175489

**Authors:** Ney Alliey-Rodriguez, Tamar A Grey, Rebecca Shafee, Jaya Padmanabhan, Neeraj Tandon, Madeline Klinger, Jonathan Spring, Lucas Coppes, Katherine Reis, Matcheri S Keshavan, Diane Gage, Steven McCarroll, Jeffrey R Bishop, Scot Hill, James L Reilly, Rebekka Lencer, Brett Clementz, Peter Buckley, Shashwath Meda, Balaji Narayanan, David C Glahn, Godfrey Pearlson, Elena I Ivleva, Carol Tamminga, John A Sweeney, David Curtis, Sarah Keedy, Judith A Badner, Chunyu Liu, Elliot S Gershon

**Affiliations:** University of Chicago, Department of Psychiatry and Behavioral Neurosciences; University of Chicago Laboratory for Advanced Computing; University of Chicago, Department of Human Genetics; Massachusetts Institute of Technology; Harvard Medical School, Department of Psychiatry; Harvard Medical School, Department of Genetics; Broad Institute of MIT and Harvard; University of Minnesota, Department of Experimental and Clinical Pharmacology; Rosalind Franklin University; Northwestern University; University of Muenster; Department of Psychology, University of Georgia, Athens; Virginia Commonwealth University; Yale University Departments of Psychiatry & Neuroscience; University of Texas Southwestern Medical Center, Department of Psychiatry; University College London and Centre for Psychiatry, Barts and the London School of Medicine and Dentistry; Rush University Medical Center; University of Illinois at Chicago

## Abstract

Schizophrenia, Schizoaffective, and Bipolar Disorders share common illness traits, intermediate phenotypes and a partially overlapping polygenic basis. We performed GWAS on deep phenotyping data, including structural MRI and DTI, clinical, and behavioral scales from 1,115 cases and controls. Significant associations were observed with two cerebrospinal fluid volumes: the temporal horn of left lateral ventricle was associated with *NRXN1*, and the volume of the cavum septum pellucidum was associated with *LRP1B* and *RORA*. Both volumes were associated with illness. Suggestive associations were observed with local gyrification indices, fractional anisotropy and age at onset. The deep phenotyping approach allowed unexpected genetic sharing to be found between phenotypes, including temporal horn of left lateral ventricle and age at onset.

## Introduction

Although very large samples of patients and controls have had genome-wide association studies (GWAS) of diagnosis of psychotic mental disorders[1], the phenotyping has been restricted. There have been very few large- or intermediate-scale studies of the many intermediate phenotypes associated with psychosis. The Bipolar and Schizophrenia Network on Intermediate Phenotypes (B-SNIP) reports here GWAS of deep phenotyping in psychosis, that is, multiple phenotypes on the same individuals, including structural magnetic resonance imaging (MRI), diffusion tensor imaging (DTI), behavioral and clinical phenotypes, from a single cohort of 1,115 individuals, including patients with schizophrenia, schizoaffective, psychotic bipolar disorder, and healthy controls. As noted by Dahl et al. [2], deep phenotyping with simultaneous genome-wide analyses serves as a discovery tool for previously unsuspected relationships of phenotypic traits with each other, and with shared molecular events. Pooling ill subjects with normal controls to analyze quantitative traits has the advantage of a wider range in the observed phenotypes, which translates into higher power to detect genetic associations[3]. All observations were generated and analyzed following standardized procedures, as discussed in prior reports from our consortium [4, 5].

The NIMH Research Domain Criteria (RDoC) initiative (https://www.nimh.nih.gov/research-priorities/rdoc/index.shtml)[6] conceives of dimensional traits that are continuous between patients and controls, and which can serve as a basis for revised definitions of disease traits and categories. In other medical disorders, such as hypertension, genetic variants have been associated with a related quantitative trait (blood pressure), once sample sizes were large enough to detect them [7]. Heritable variance in blood pressure and clinical hypertension are accounted for by GWAS-significant associations with common variants, by polygenic risk, and by rare variants with strong effect[8], which is generally true for common diseases[9].

There is ample evidence of partially overlapping brain morphology abnormalities in schizophrenia (SZ) and bipolar disorder (BD) [10-12]. These disorders also have a shared set of specific genetic loci associated with them, and a partially shared polygenic diathesis [1, 13], as well as shared behavioral features including psychosis in some BD patients. Identifying genetic factors associated with specific biological traits associated with psychosis in these disorders can help to elucidate the biology of shared disease features, which may lead to improved disease classification and personalized treatment [14].

Previous studies have identified enlargement of the ventricular system associated with bipolar disorder and schizophrenia [15, 16]. The temporal (inferior) horn of the left lateral ventricle and the cavum septum pellucidum (CSP) are two structures that have consistently been found enlarged in patients with these disorders [17-19]. In this study we found common variants of genes Neurexin1 (*NRXN1*), LDL receptor related protein 1B (*LRP1B*), and RAR-Related Orphan Receptor (*RORA*) associated with enlargement of these structures. A previous study by Rujescu et al. described rare structural genetic variants of *NRXN1* associated with Schizophrenia [20]. *NRXN1* encodes a large presynaptic transmembrane protein that binds neuroligins to calcium-dependent synaptic complexes in the central nervous system, and is also involved in the formation of synaptic contacts [21]. *LRP1B* is a candidate gene for Alzheimer’s disease [22] and *RORA* is a candidate gene for PTSD [23].

## Results

We identified 31 SNP markers with at least a suggestive genome-wide association (P < 1E-08) in 21 out of 463 total studied phenotypes in the B-SNIP first wave (B-SNIP1) sample, which contains 1,115 unrelated cases and controls genotyped using the Illumina Infinium Psycharray (PsychChip) followed by imputation to the 1000 Genomes reference panel, and filtered to work with common SNPs (MAF≥0.05) on the PLINK regression model GWAS. There were three significant results after Family-Wise Error Rate (FWER) correction (see Methods) in the CSF volumes phenotype group. The strict Bonferroni significance threshold after multiple testing for 463 phenotypes is P < 2.16E-11 which was exceeded by the same three top results. Here we tabulate results with the P < 1E-08 suggestive association threshold, but only discuss significant results. Complete GWAS results can be requested from the authors through the B-SNIP website (http://b-snip.org).

**Table 1:**
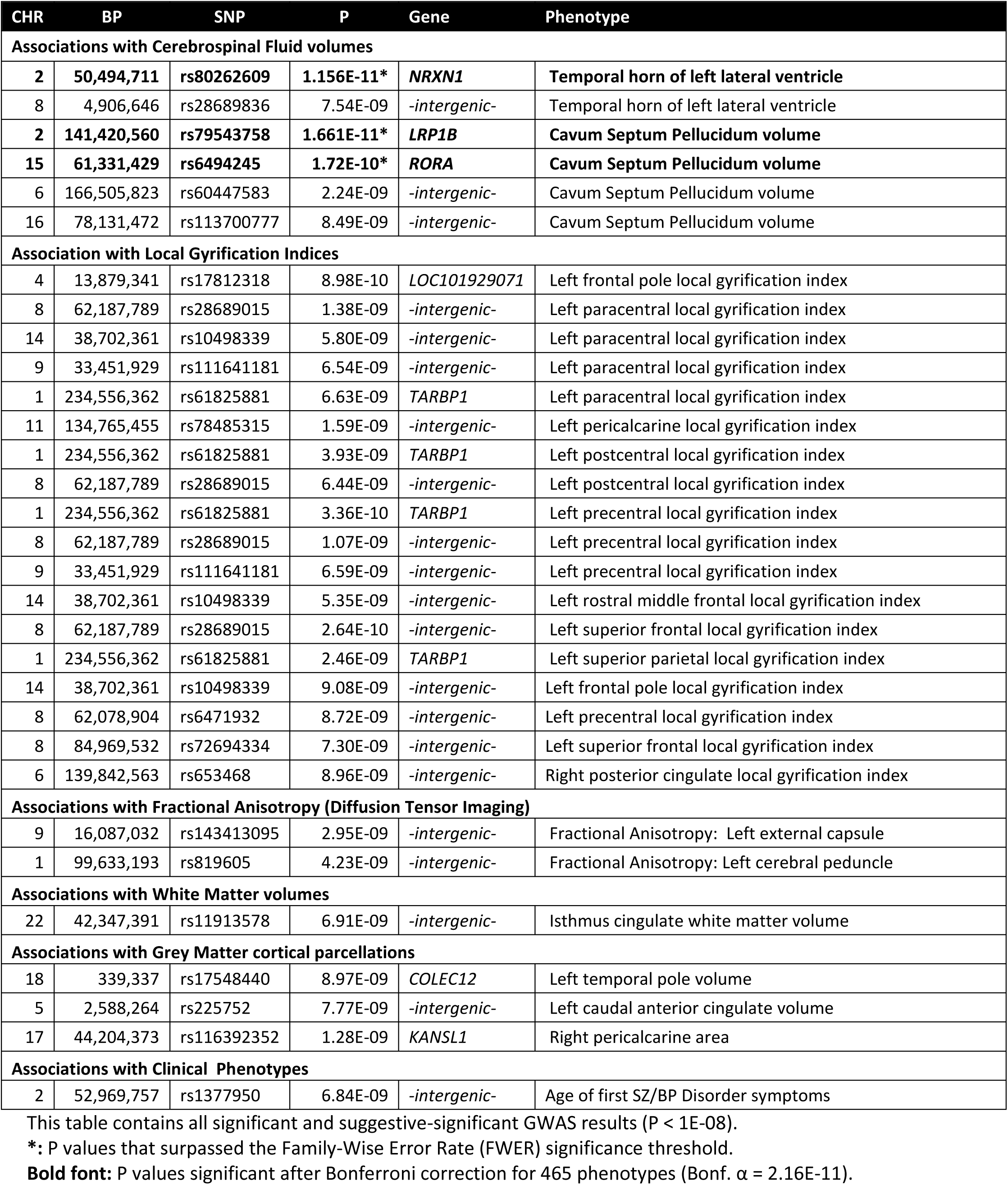
Genome-Wide Association results:

### Associations with Cerebrospinal Fluid volumes

The most significant genome-wide P values observed were associated with two CSF volumetric phenotypes measured by FreeSurfer structural MRI: the volume of the temporal horn of left lateral ventricle was associated (P= 1.156E-11) with rs80262609, an intronic SNP of Neurexin1 gene (*NRXN1*) located at Chr.2p16.3 (Fig.1a); and the volume of the Cavum Septum Pellucidum (CSP) (Fig.1b) had P= 1.661E-11 in association with rs79543758, an intronic SNP of the LDL receptor-related protein 1B gene (*LRP1B*). This same phenotype also had a P= 1.72E-10 association with rs6494245, an intronic SNP of the RAR-Related Orphan Receptor A gene (*RORA*) (Fig.1.b.2), which is a PTSD candidate gene [23]. These two CSF volumetric variables showed significant differences between cases and controls in the studied sample: The temporal horn of left lateral ventricle (ANOVA P=0.01) and the CSP (ANOVA P=9.4E-05) were larger in cases (Figure 2). A post-hoc analysis using Quanto [24] calculated the power for these two top SNP findings as 0.62 and 0.57, and their effect sizes (Eta) were 0.249 and 0.265 respectively (Supplementary Table 4). Because of reported discrepancies in reports on the association of CSP with SZ [25, 26], we conducted a blind semi-quantitative verification of the Freesurfer results with a random selection of MRI images by experienced imaging scientists using manual measurements, obtaining a positive correlation between manual ratings and the FreeSurfer CSP quantitative output (FreeSurfer vs. Rater’s Cronbach’s Alpha=0.775 (P=4.37E-07).

**Figure 1:**
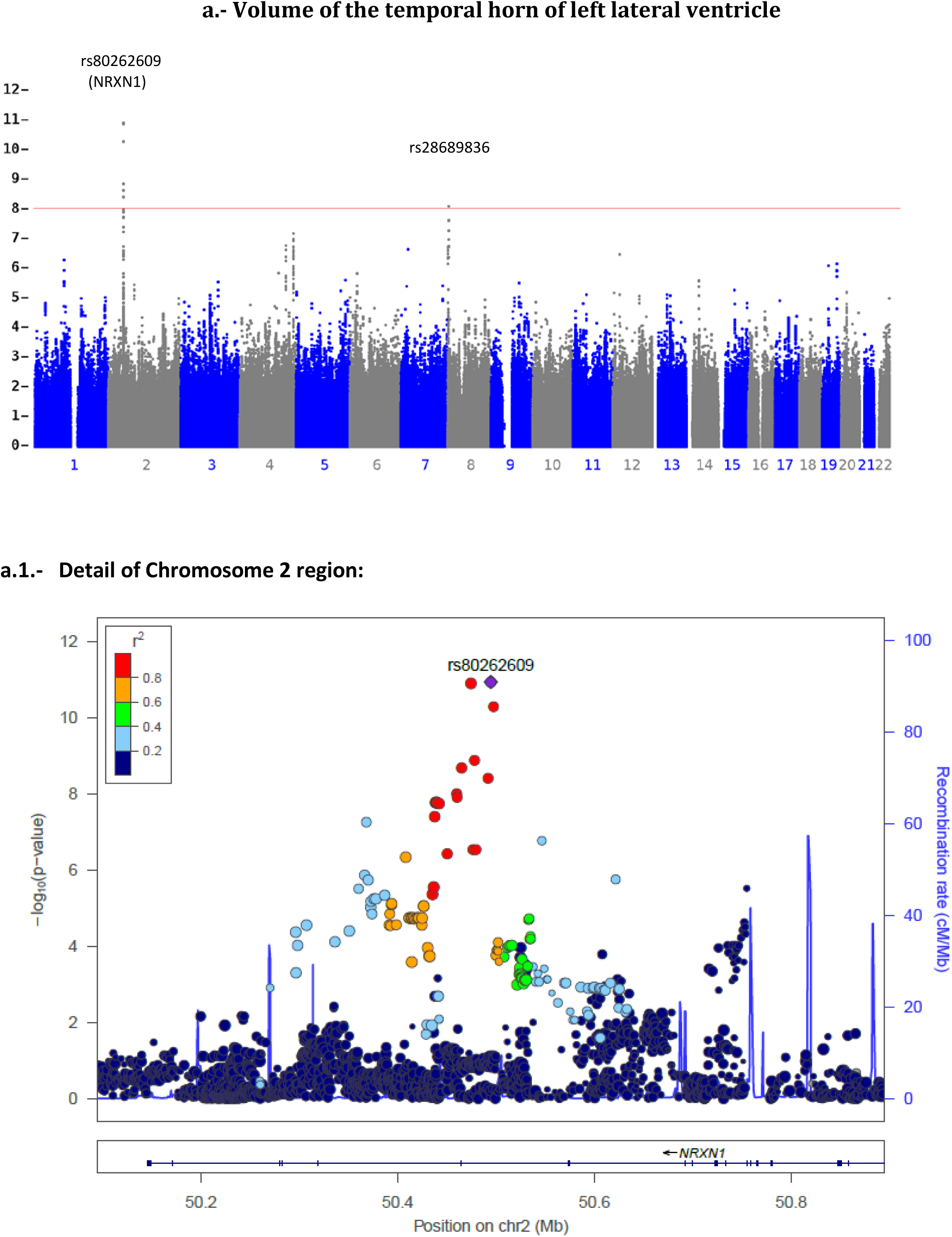
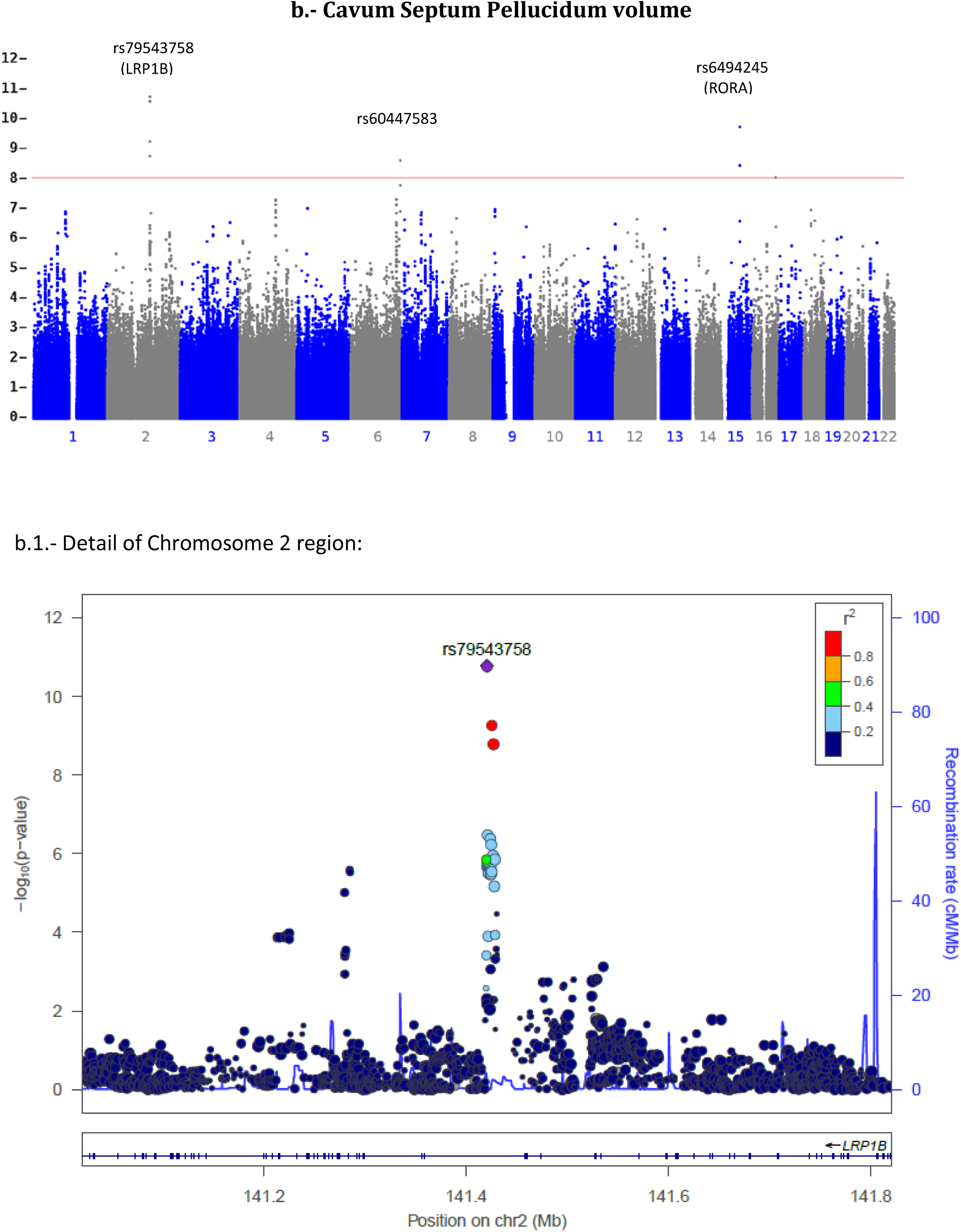
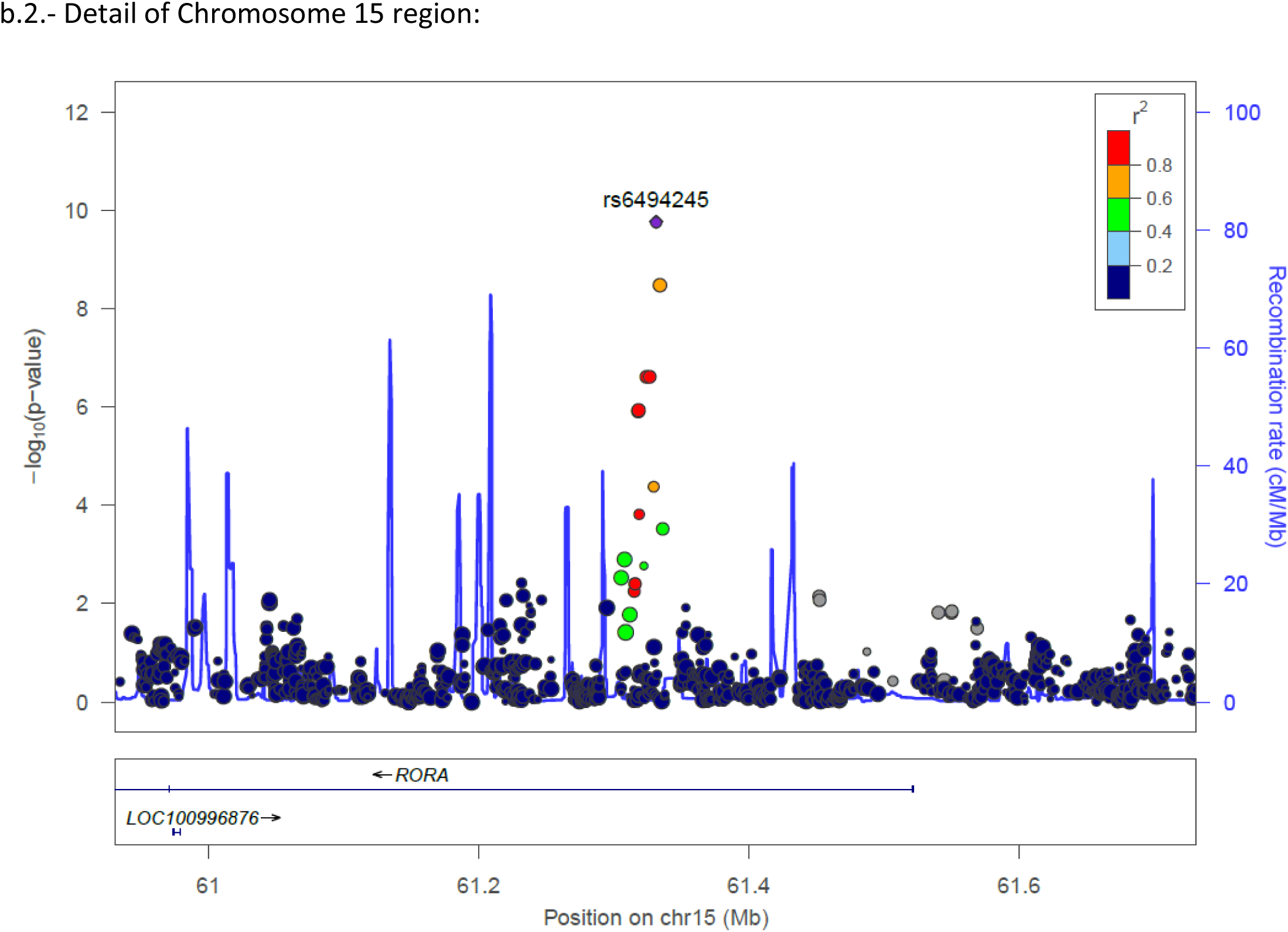
Manhattan plots of top GWAS results:

**Figure 2:**
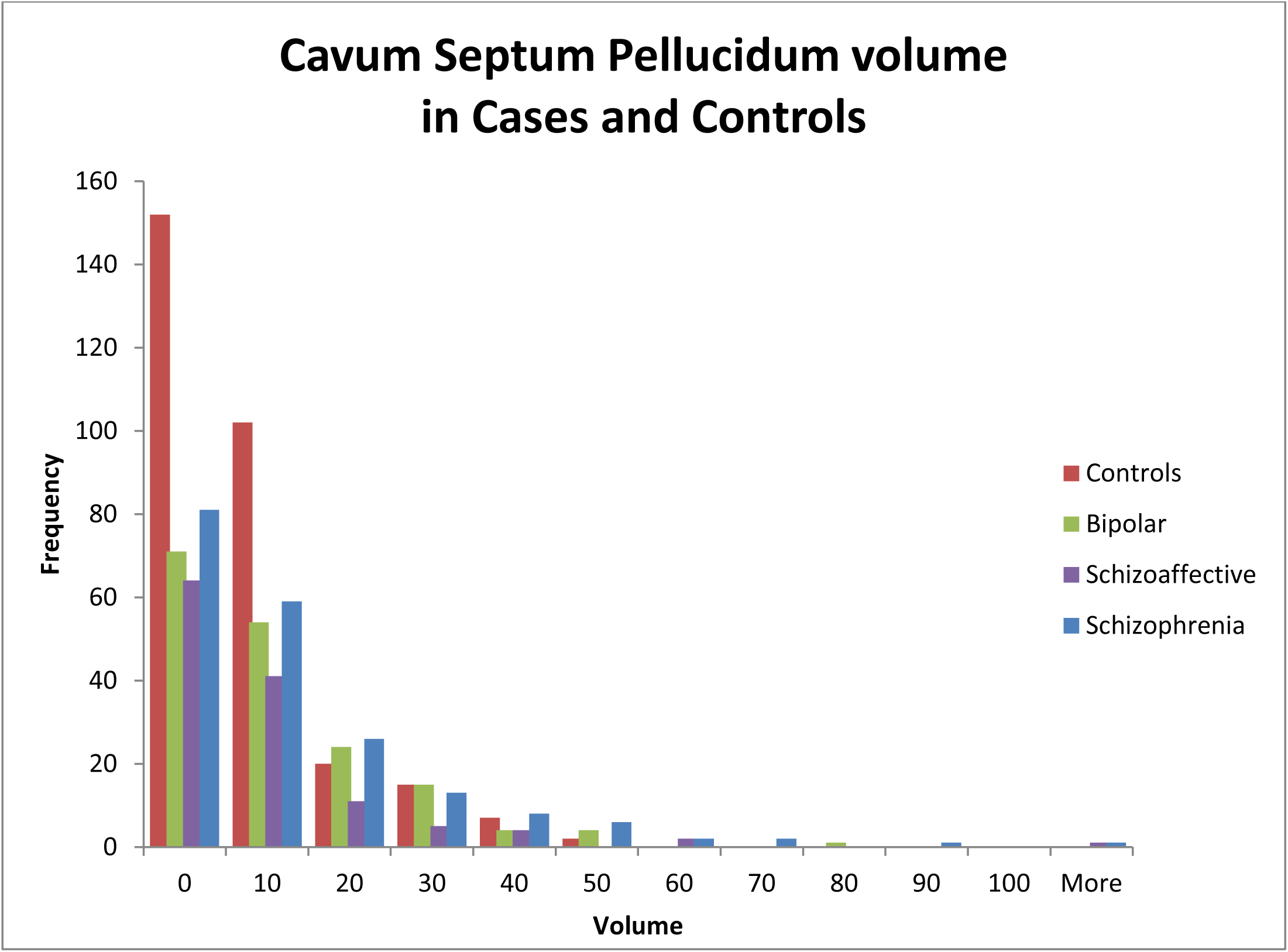
Cavum Septum Pellucidum volume distribution in Cases and Controls. The distribution of CSP volumes (in Voxels) in cases and controls by diagnostic categories identified significant differences between groups (ANOVA P=9.4E-05).

### Other suggestive GWAS results

### Associations with Local Gyrification Indices

FreeSurfer-based Local Gyrification Indices (LGI) from 9 brain regions showed 13 suggestively significant associations. Four of them (left precentral, left postcentral, left paracentral, and left superior parietal LGIs) were suggestively associated with the Homo sapiens TAR (HIV-1) RNA binding protein 1 gene (*TARBP1*). Three of these LGIs and one additional region (left precentral, left postcentral, left paracentral and left superior frontal LGIs) had suggestive association with rs28689015, an intergenic SNP on Chr8q12.2; and the left frontal pole LGI with rs17812318, an intronic SNP of *LINC01182*, a long non-coding RNA. Four other LGI suggestive associations pointed to other intergenic regions with P<1E-08 (Table 1). Analysis of variance did not show significant differences between cases and controls or among diagnostic categories for these phenotypes.

### Associations with Fractional Anisotropy

Fractional Anisotropy (FA) is a metric of white matter fiber density and integrity, computed from Diffusion Tensor Imaging (DTI). FA of the left external capsule was suggestively associated (P=2.947-E-09) with rs143413095, an intergenic SNP on Chr.9p23.3; and FA of the left cerebral peduncle had suggestively significant association (P=4.233E-09) with rs819605, an intergenic SNP on Chr.1p21.3. No significant differences between cases and controls were observed for these two variables.

### Association with Age at Onset

One clinical phenotype, the “Age of first symptoms of schizophrenia or bipolar disorder” reached suggestive significance association with rs1377950 (P=6.841E-09), however this did not pass the FWER significance threshold for psychiatric and medical history variables (Supplementary Table 1). This SNP is an intergenic marker on Chr.2p16.2, 300Kb from *LOC730100*, a long non-coding RNA neighbor of the *NRXN1* gene. A microRNA (*Mir4431*) has also been described adjacent to this locus. The CSP volume had a small but significant negative correlation with the age at onset in our sample (R= -0.118; P=3.89E-04).

## Discussion

### Significant genetic associations with brain CSF volumetric variants related to psychotic disorders

Cerebrospinal fluid volumes are some of the most reliable measurements captured with *in vivo* MRI, because of the high contrast between CSF and cerebral structures. Enlarged ventricular volumes have been among the most enduring anatomical findings in SZ and BD, first reported by Johnstone in schizophrenia forty years ago [27]. This has been confirmed in SZ and BD patients compared with healthy controls in large studies by the ENIGMA consortium [16, 28], and its heritability calculated as 0.54[29]. Increased ventricular volumes observed in schizophrenia have been attributed to volume reduction of surrounding grey matter structures [30, 31]. The temporal horn of the lateral ventricle traverses the temporal lobe and is in direct relation with the amygdala, hippocampus, and caudate nucleus. However, our GWAS on these structures’ separate volumes did not have significant associations, which implies that the volumetric association may reflect joint regulation of part of the morphology of these structures. We noted that hippocampal volumes in our sample were significantly smaller in cases than in controls (ANOVA P=3E-06)[32-34], and there were also significant reductions in cases for the amygdala volumes (P=0.001), but not for the caudate. Gaser et al. reported ventricular enlargement in schizophrenia associated with volumetric reduction of the thalamus and superior temporal cortex [30]; in our data, however, we did not observe significant correlation between the volumes of these structures and the volume of the temporal horn of lateral ventricle, or any detectable associations between these structures volumes and the CSF volume-associated genes.

In our analyses we observed significantly larger volumes of the temporal horn of left lateral ventricle in cases vs. controls, and polymorphism in the Neurexin1 gene (*NRXN1*) was associated with this endophenotype. When we combined temporal horns values across hemispheres, the association was still suggestively significant (P=4.92E-10). The Psychiatric Genomic Consortium (PGC) reported an association P= 0.005685 for this same SNP marker of *NRXN1* (rs80262609) with diagnosis in their large case-control schizophrenia GWAS [1]. Structural variants (CNVs) disrupting this gene have been associated with a wide spectrum of brain disorders including schizophrenia [20, 35, 36], and with cognitive performance through the regulation of post-synaptic N-methyl d-aspartate receptors [37]. This gene is expressed in the adult brain[38], and during development in the fetal brain[39].

The cavum septum pellucidum is a space between the two leaflets of the septum pellucidum that normally occludes in 85% of babies by age 6 months[40]; its persistence has been related to neurodevelopmental delay[41], and with SZ and BD[17, 18, 42, 43], presumably because of a reduction of the limbic structures that surround it: the septal nuclei, fornix and hippocampal commissure. In our sample this quantitative phenotype measured in voxels by FreeSurfer was significantly different in cases vs. controls (ANOVA P=9.4E-05, Figure 2), with larger volumes for individuals with SZ than BP, which is consistent with findings from Shioiri et al. [18]. One of the genes identified in association with the CSP volume, the LDL receptor related protein 1B, has been associated with Attention Deficit/Hyperactivity Disorder and Alzheimer’s disease [22, 44]. *LRP1B* is reported expressed in the adult and fetal brain[45] which, along with its association with an anatomical quantitative trait (CSP) of a persistent fetal structure, supports a role of this gene in a neurodevelopmental hypothesis of Bipolar Disorder and Schizophrenia [26]. *RORA* is a nuclear receptor that participates in the transcriptional regulation of some genes involved in circadian rhythm and in the response to lithium treatment in bipolar disorder[46]. In mice, RORα is essential for development of cerebellum through direct regulation of genes expressed in Purkinje cells.

According to UK Brain Expression Consortium (UKBEC) eQTL data (http://www.braineac.org/), rs80262609 is an eQTL SNP (P < 0.01) for *NRXN1* in putamen area, temporal cortex, frontal cortex, and thalamus (Supplementary Table 3), whereas rs6494245 is primarily an eQTL SNP for *VPS13C* in hippocampus, cerebellum, and occipital cortex; also associated with expression of *RORA* in hippocampus, frontal cortex, medulla, and thalamus; associated with *NARG2* expression in hippocampus, cerebellum, and frontal cortex; with *ANXA2* expression in hippocampus. Among these genes, *RORA* is a core transcription factor involving circadian rhythms. *VPS13C* involves mitochondrial function and maintenance of mitochondrial transmembrane potential. *NARG2* regulates small nuclear RNA (snRNA) gene transcription. *ANXA2* involves in exocytosis. Co-expression analysis of *NRXN1* and *LRP1B* using the Neuropsychiatric Disorder De Novo Mutations Database [47] identified genes involved in cell adhesion (including Cadherin13) and neurotransmission (including the Glutamate Ionotropic Receptor AMPA 2 (Supplementary Figure 3). Previously published B-SNIP analyses on a subset of this sample looking for gene networks associated with some of these phenotypes found cell adhesion networks containing CDH13, NRXN3 and *RORA* in association with EEG oscillations, event-related potentials, resting state fMRI and structural MRI phenotypes [12, 48-51]. Although not a replication because our data contains that subsample, this study may represent support for these previous studies with respect to CDH13 and *RORA*.

The significant genetic associations with volumes of the temporal horn of lateral ventricle and of the CSP represent novel genetic findings on the molecular basis of psychotic disorders, and are the first genetic associations with psychosis that pinpoint a stage of brain development by direct anatomic observation. The relationship between age of onset of psychosis and a ventricular volume, in the context of a significant genetic association for one of the two phenotypes and a suggestive association for the other, is the type of finding that gives credence to the B-SNIP consortium deep phenotyping approach study [2]. We have observed highly significant p-values with a relatively smaller sample size than needed for disease association, and relatively large effect sizes for the significant associations, which suggest that diagnosis may be a more difficult phenotype to study than some of its observable components.

## Methods

### The BSNIP1 data set

Data analyzed here comes from the Bipolar and Schizophrenia Network for Intermediate Phenotypes - first wave (B-SNIP1), which studied individuals with a history of psychosis and diagnoses of Schizophrenia, Bipolar, or Schizoaffective disorder (SAD) [52]. BSNIP-1 collected a comprehensive battery of phenotypes including clinical and behavioral interview data, cognitive trait assessments, oculo-motor testing with smooth pursuit and saccade paradigms, resting-state EEG, auditory-event-related potentials, and brain imaging (structural, diffusion tensor, resting state functional MRI). This comprehensive set of psychosis-related phenotypes was acquired between 2008 and 2012 from more than 2,400 individuals, including 312 BD with psychotic features, 223 SAD and 397 SZ cases, 1029 first degree relatives of cases, and 455 unrelated healthy controls following standardized methods in five different study sites in the USA, under IRB approval from each participating institution. Relatives were excluded from our GWAS analyses and only individuals that passed QC described below were included.

### Phenotypes

Details of B-SNIP1 phenotype collection are described in Tamminga el al. 2014 [5], Diffusion Tensor Imaging was processed as in Skudlarski 2010[11], and structural neuroimaging variables extracted from MRI were processed using FreeSurfer version 5.1 [53] as described in Padmanabhan et al. 2015[10]. These phenotypes, assembled within the B-SNIP1 analytical database [5], were used as quantitative phenotypes for genome-wide association. Analyses were conducted on data from 1,115 unrelated patients and healthy control volunteers with available phenotypes and genotypes: of these 754 are cases (245 Bipolar, 186 Schizoaffective and 323 Schizophrenia) and 361 controls; 561 are males and 554 are females. 654 individuals self-reported as Caucasians, 375 as African Americans, and 86 self-reported other ethnicities (includes Asians, American Indian, Native Hawaiian, multiracial, others and unknown). The number of subjects analyzed for each trait varied, as only individuals with non-missing data were included for analysis and no phenotype imputation was applied. Phenotypes were inspected for accuracy and to detect outliers based on their values distribution. A total of 463 phenotypes were used for GWAS. Supplementary Table 1 shows a summary of phenotypes by groups, and the complete phenotype list is available in Supplementary Table 2. Although some variables exhibited high correlation with others like thickness, area and volumes from structural MRI cortical parcellations, genetic factors associated with each may be different and worth analyzing separately as stated by Winkler et al. [54].

### Genotypes

were assessed from blood DNA at the Broad Institute using the Illumina Infinium Psycharray (PsychChip), which contains a total of 588,454 SNP markers, including 50,000 specific genetic markers for neuropsychiatric disorders. PsychChip genotype calls were processed through a Broad Institute custom pipeline designed to merge calls from 3 different algorithms (GenCall, Birdseed and zCall), in order to maximize reliability and usability of rare markers [55]. Pre-Imputation QC applied to the called PsychChip genotypes included filtering by Call Rate > 98% by SNP and >98% by sample, HWE P-value > 1E-06 in controls, Inbreeding Coefficient (-0.2 > F_Het > 0.2), exclusion of monomorphic markers, Sex Check using X chromosome heterozygosity and Y chromosome call rate, and Minor Allele Frequency (MAF) 0.01. We used programs PREST-plus [56] and KING [57] to check genotypes of all putatively unrelated patient and control individuals for cryptic relatedness, so that the final dataset had no individuals who showed 3^rd^ degree or closer kinship.

### Genotype Imputation

After QC, PsychChip genotypes were imputed to the 1000 Genomes project multiethnic reference panel [58] using HAPI-UR for pre-phasing and IMPUTE2 [59-61]. Chromosomes were phased separately, and then divided into 5 megabase pair chunks for imputation. Poorly imputed single nucleotide polymorphisms (SNPs) were filtered post-imputation. The final imputed genotype set contained more than 30 Million autosomal markers, reduced to 4,322,238 variants after filtering for 0.05 missingness by marker, 0.02 missingness by individual sample and MAF 0.05, to work with common variants.

### GWAS analysis

were performed using PLINK 1.9 [62, 63] on the Bionimbus protected cloud of the Open Science Data Cloud servers at the University of Chicago [64](http://www.opensciencedatacloud.org). Genetic locations refer to the human genome GRCh37/hg19 build, gene mapping was done using the UCSC genome browser [65], and graphics generated with Manhattan Plotter[66] and LocusZoom[67]. Population stratification was expected in the B-SNIP1 sample, as it contains people from diverse ethnicities. We ran Principal Component Analysis (PCA) on the genotypes to examine population stratification and admixture in our sample (Supplementary Figure 1a). Then, the first two PCA eigenvectors which captured the vast majority of the ethnic-related variance (Supplementary Figure 1b) were used as covariates in the GWAS regression model to correct for population stratification and admixture [68]. Sex was also used as a covariate in all our analyses. Although recent publications use multiple additional covariates (like case-control status, age, total intracranial volume, etc.) [16, 69], in our study we explicitly avoided them because correcting for factors that are related to the disease could diminish the possibility of detecting associations with loci that differentiate cases and controls, or that are related to the evolution of the disease and that can thus be responsible for phenotypic variance. Where significant or suggestive gene associations were observed with a phenotype, analysis of variance (ANOVA) was used to test for significant case-control phenotypic differences, calculated with IBM SPSS Statistics version 24. Q-Q plots were used to examine possible inflation in GWAS results (Suppl. Figure 2).

### Significance Thresholds

The classic definition of statistically suggestive and significant probability thresholds in whole genome analysis is that a result is suggestive if it is expected to occur not more than once in a single genome-wide analysis, and significant if it is expected to occur not more than once in the context of all the analyses under consideration[70]. The GWAS statistical significance threshold for a single phenotype analysis of imputed data using the 1000 Genomes reference panel is accepted as P < 1E-08, based on the number of effective markers (Li et al. [71]). In the context of the multiple analyses reported here, this would be our threshold for suggestive significance. Given the different classes of phenotype classes analyzed (psychiatric scales, clinical data, fractional anisotropy and structural MRI, we opted for a Family-Wise Error Rate (FWER) approach (considering each class of variable as a separate family) to correct for multiple testing and declare FWER significance (Supplementary Table 1). We note separately findings that pass a Bonferroni correction for all variables analyzed.

## Acknowledgments

Thanks to the probands, their families and volunteers who joined this study and contributed their time and individual data.

This work made use of the Bionimbus Protected Data Cloud (PDC), which is a collaboration between the University of Chicago Center for Data Intensive Science (CDIS) and the Open Commons Consortium (OCC). The Bionimbus PDC allows users authorized by NIH to compute over human genomic data in a secure compliant fashion. The Bionimbus PDC is part of the OSDC ecosystem.

## SUPPLEMENTARY MATERIAL

**Supplementary Table 1:** Family-Wise Error Rate significance threshold by phenotype groups. Complete list of phenotypes included in each group is available in Supplementary Table 2.

Family-Wise Error Rate significance threshold by phenotype groups
| Phenotype Groups | Number of Phenotypes | FWER $\alpha$ | FWER Significant Results |
| --- | --- | --- | --- |
| **1.- Psychiatric Scales** | **15** | 6.25E-10 | - |
| **2.- Psychiatric and Medical History** | **4** | 2E-09 | - |
| **3.- Fractional Anisotropy, DTI** | **50** | 2E-10 | - |
| **4- FreeSurfer structural MRI** | **377** | 2.65E-11 |  |
| CSF volumes | 7 | 1.43E-09 | 3 |
| Cortical parcellations LOCAL GYRIFICATION INDICES | 68 | 1.47E-10 | - |
| Grey Matter Cortical parcellations AREAS | 70 | 1.43E-10 | - |
| Grey Matter Cortical parcellations THICKNESS | 68 | 1.47E-10 | - |
| Grey Matter Cortical parcellations VOLUMES | 68 | 1.47E-10 | - |
| Sub-Cortical Structures AREAS | 2 | 5E-09 | - |
| Sub-Cortical Structures VOLUMES | 12 | 8.33E-10 | - |
| White Matter Cortical parcellations VOLUMES | 68 | 1.47E-10 | - |
| White Matter | 6 | 1.67E-09 | - |
| Cerebellum | 4 | 2.5E-09 | - |
| Others | 4 | 2.5E-09 | - |
| **Additional FreeSurfer global variables:** | **17** |  |  |
| Total CSF volume | 1 | 1E-08 | - |
| Grey Matter | 5 | 2E-09 | - |
| Large brain structures | 6 | 1.67E-09 | - |
| White Matter | 5 | 2E-09 | - |
| **All Phenotypes (Bonferroni)** | **463** | **2.16E-11** | **3** |

**Supplementary Table 2: Phenotype list**

Download it from: https://drive.google.com/file/d/0B_abC_nlCXwHMnl6SThsWXYwRG8/view?usp=sharing

**Supplementary Table 3: eQTL analysis**

Download it from: https://drive.google.com/file/d/0B_abC_nlCXwHc29FdHlGZGpKVGc/view?usp=sharing

**Supplementary Table 4:** Power Analysis for top SNP findings. Power analysis for top SNPs in BSNIP GWAS using significance threshold = 1.78E-11, calculated with SPSS, Quanto and verified with G*Power[72].

| Phenotype | Marker | N | MAF | Effect Size | Power |
| --- | --- | --- | --- | --- | --- |
| Temp Horn Left Lat Ventr | rs80262609 | 789 | 0.0538 | 0.249 | 0.62 |
| CSP | rs79543758 | 784 | 0.0554 | 0.265 | 0.57 |

**Supplementary Figure 1a:**
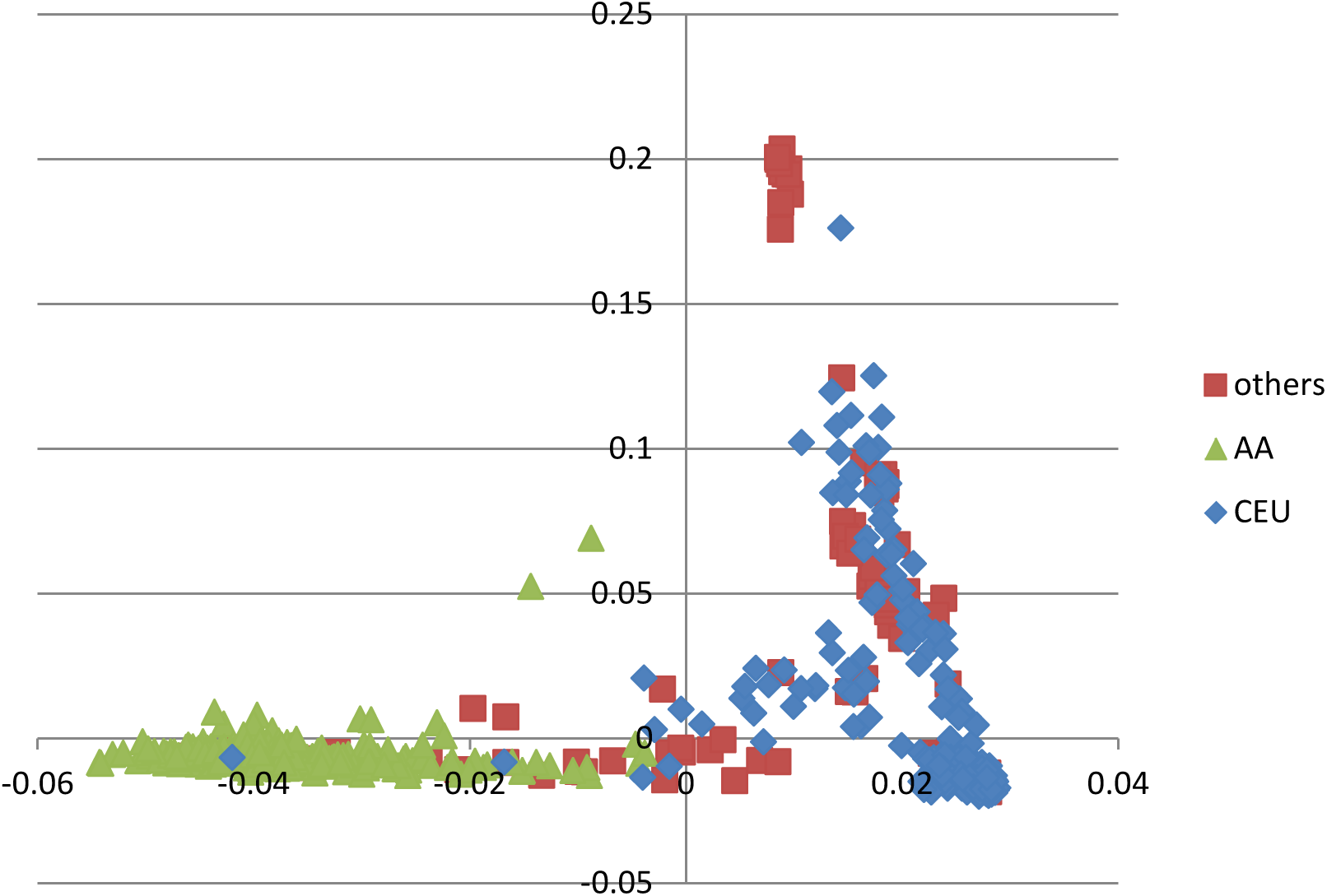
Principal Component Analysis (PCA) of genotypes showing population stratification. *Genotype’s PCA vs. Self-Reported ethnicity: Plot of the 2 first Eigenvectors from Principal Component Analysis (PCA) of genotypes: lower left corner cluster corresponds to African ancestry (AA), lower right corner cluster to Caucasian ancestry (CEU), and the upper middle to Asian ancestry component. Color code is self-reported ethnicity.*

**Supplementary Figure 1b:**
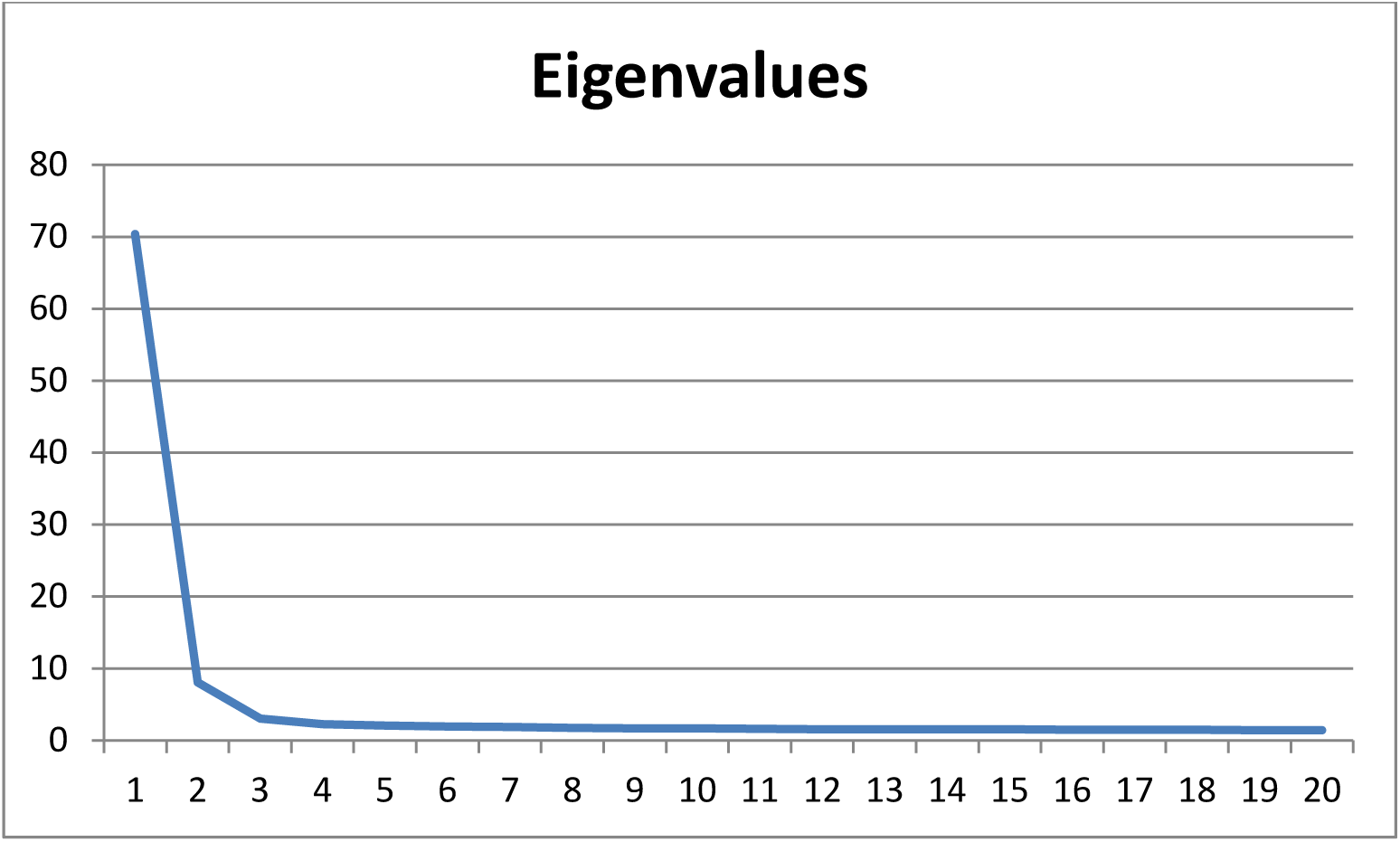
Plot of Eigenvalues from Genotypes’ PCA, showing the variance captured by each Eigenvector. The two first Eigenvectors, which captured the majority of ethnic-related variance were used as covariates for GWAS.

**Supplementary Figure 2:**
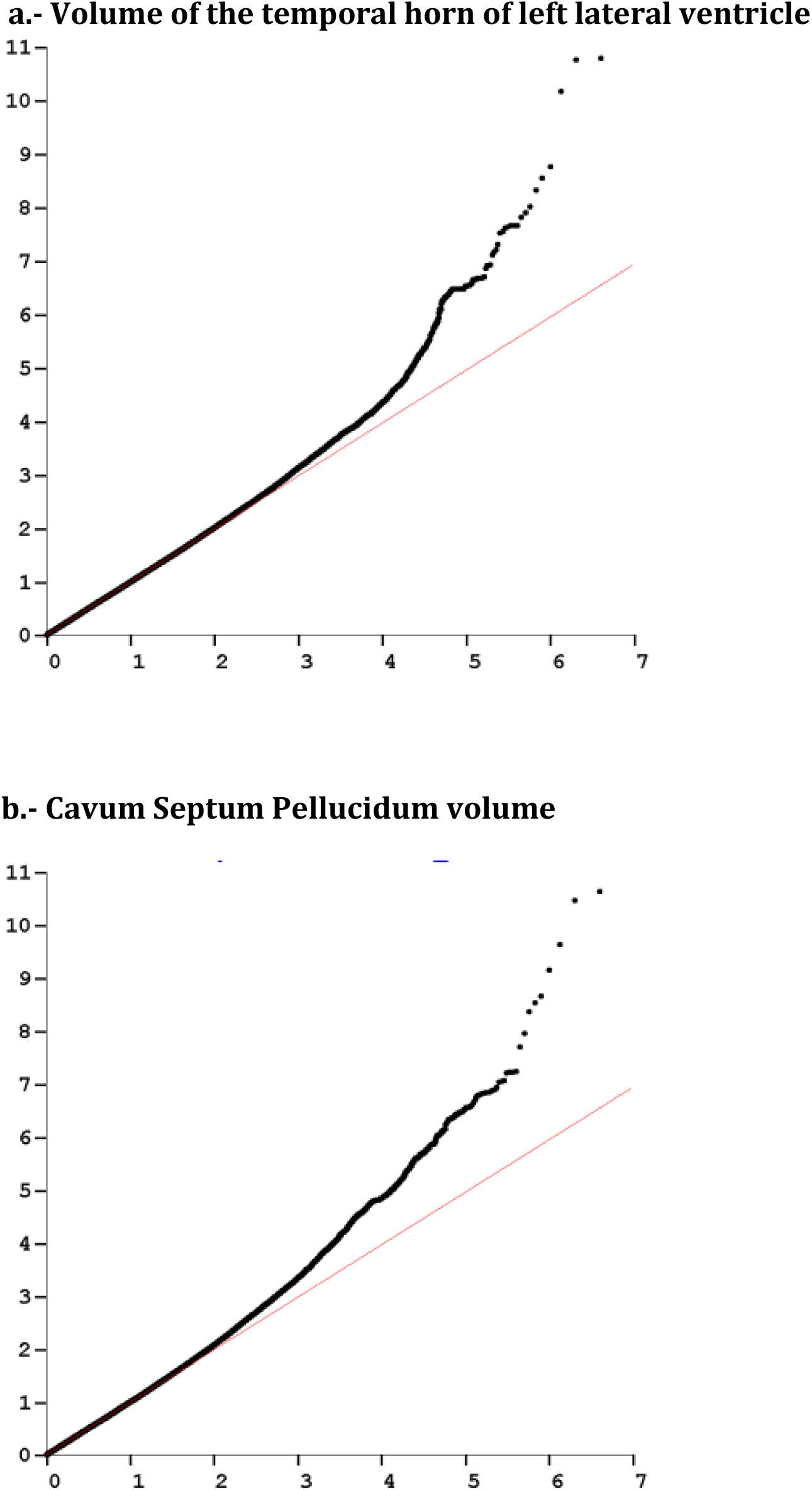
GWAS Quantile-Quantile plots for highlighted phenotypes. The red line represents the null distribution

**Supplementary Figure 3:**
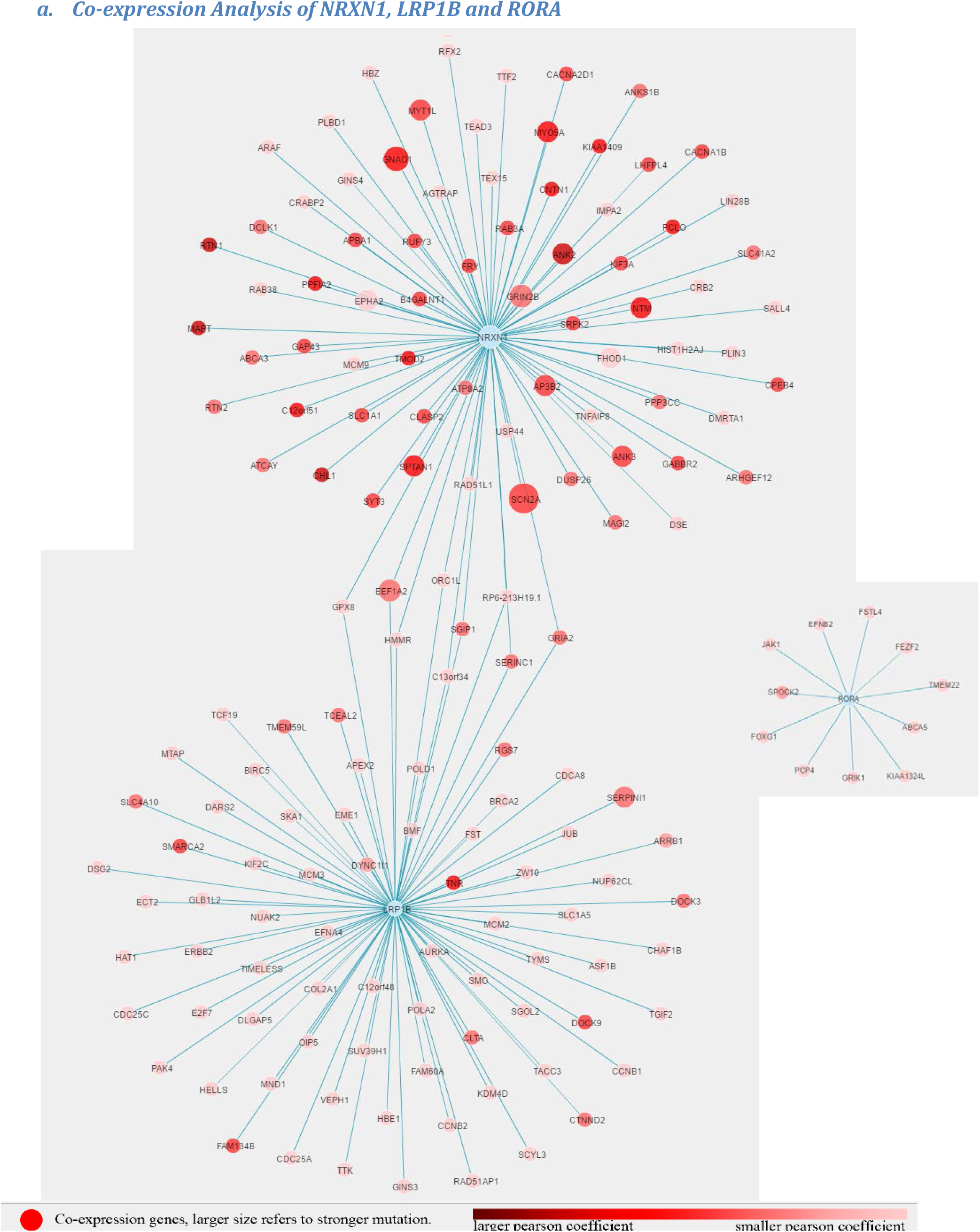
a. Co-expression Analysis of NRXN1, LRP1B and RORA. Co-expression analysis of the 3 genes associated with ventricular enlargement (NRXN1, LRP1B and RORA), using NPdenovo Co-expression analysis (http://www.wzgenomics.cn/NPdenovo/index.php). Reference Dataset: HBT; Minimum Pearson correlation coefficient: 0.8; Maximum nodes of network: 80. Several genes are co-expressed by NRXN1 and LRP1B, including GRIA2. No co-expression with RORA.

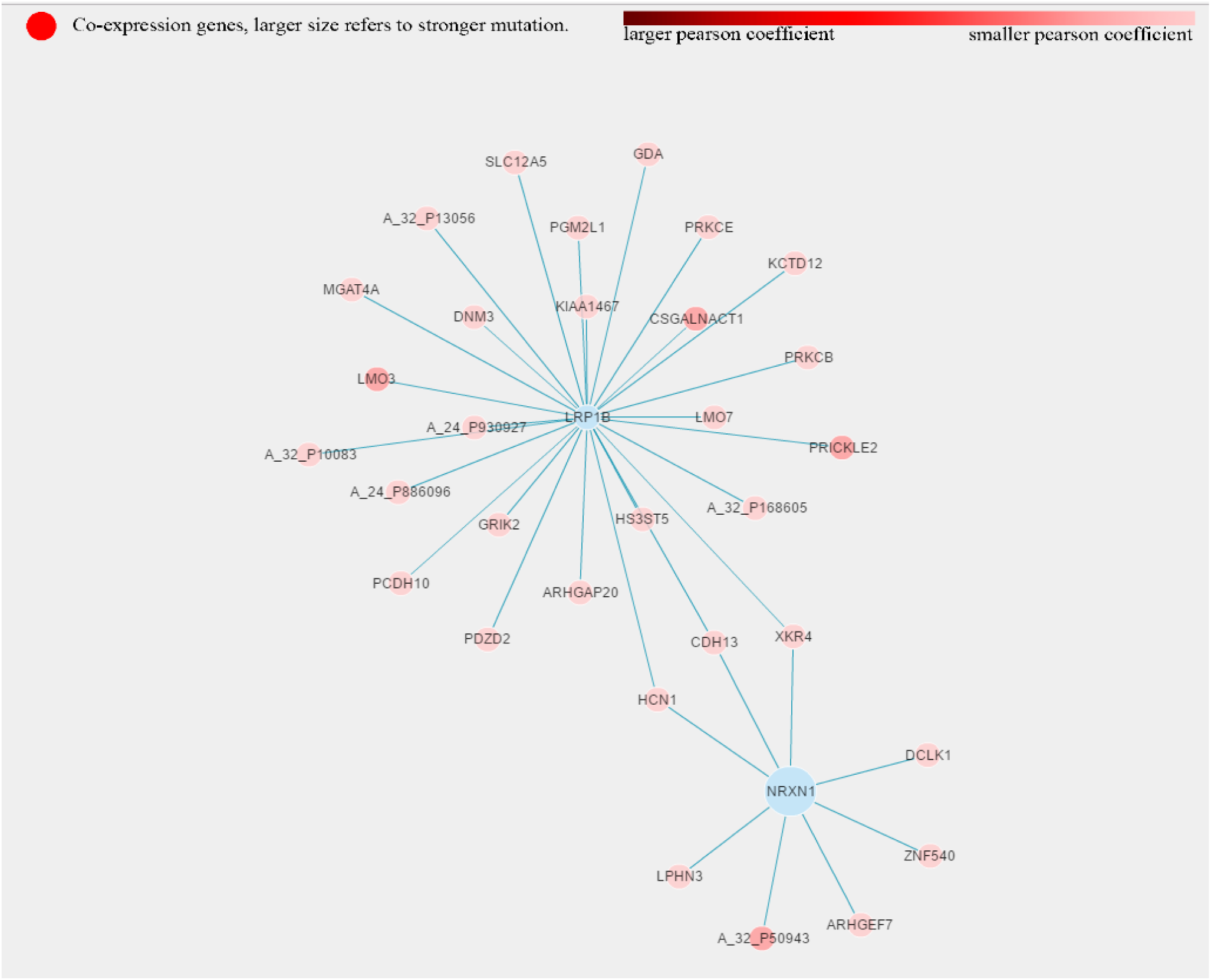
b. Additional Co-expression Analysis of NRXN1 and LRP1B. Secondary Co-expression analysis of NRXN1 and LRP1B on NPdenovo Co-expression analysis (http://www.wzgenomics.cn/NPdenovo/index.php), using Dataset: LMD, Minimum Pearson correlation coefficient: 0.8 Maximum nodes of network: 80. Cadherin 13 (along with 2 other genes) is identified in this dataset as a co-expressed gene.

